# Serotonergic regulation of corticoamygdalar neurons in the mouse prelimbic cortex

**DOI:** 10.1101/293688

**Authors:** Daniel Avesar, Emily K. Stephens, Allan T. Gulledge

**Affiliations:** Department of Molecular and Systems Biology Geisel School of Medicine at Dartmouth College 74 College St., Vail 601 Hanover, New Hampshire, 03755 USA

## Abstract

Neuromodulatory transmitters, such as serotonin (5-HT), selectively regulate the excitability of subpopulations of cortical projection neurons to gate cortical output to specific target regions. For instance, in the mouse prelimbic cortex, 5-HT selectively excites commissurally projecting intratelencephalic (COM) neurons via activation of 5-HT_2A_ (2A) receptors, while simultaneously inhibiting corticofugally projecting pyramidal neurons targeting the pons via 5-HT_1A_ (1A) receptors. Here we characterize the physiology, morphology, and serotonergic regulation of corticoamygdalar (CAm) projection neurons in the mouse prelimbic cortex. Layer 5 CAm neurons shared a number of physiological and morphological characteristics with COM neurons, including higher input resistances, smaller HCN-channel mediated responses, and sparser dendritic arbors than corticopontine neurons. Across cortical lamina, CAm neurons also resembled COM neurons in their serotonergic modulation; focally applied 5-HT (100 µM; 1 s) generated 2A-receptor-mediated excitation, or 1A- and 2A- dependent biphasic responses, in ipsilaterally and contralaterally projecting CAm neurons. Serotonergic excitation depended on extrinsic excitatory drive, as 5-HT failed to depolarize CAm neurons from rest, but could enhance the number of action potentials generated by simulated barrages of synaptic input. Finally, using dual tracer injections, we identified double-labeled CAm/COM neurons that displayed primarily excitatory or biphasic responses to 5-HT. Overall, our findings reveal that prelimbic CAm neurons overlap with COM neurons, and that both neuron subtypes exhibit 2A-dependent serotonergic excitation. Our findings suggest that 5-HT, acting at 2A receptors, will promote cortical output to the amygdala.

**Significance Statement:** Cortical projections to the amygdala allow for executive “top-down” control of emotional responses. Corticoamygdalar (CAm) neurons in the prelimbic cortex contribute to the learning and expression of conditioned fear responses, processes that may also be regulated by serotonin (5-HT). Our study provides a physiological and morphological characterization of prelimbic CAm neurons, and demonstrates that the excitability of CAm neurons is regulated by 5-HT acting at 5-HT_2A_ receptors alone, or in combination with 5-HT_1A_ receptors. Our results suggest that 5-HT may regulate corticoamygdalar circuits during the learning and expression of conditioned fear.

## Introduction

Output from the neocortex is organized into multiple information channels formed by subclasses of glutamatergic pyramidal neurons defined by their long-distance axonal projections to distinct cortical and subcortical targets. Cortical projection neurons are differentially regulated by a variety of neuromodulatory transmitters through cell-type-specific expression of receptors, signaling proteins, and ionic effectors. For instance, in the mouse prelimbic cortex, acetylcholine (ACh) and serotonin (5-HT) have opposing influences on the excitability of two broad classes of projection neurons: corticofugal neurons, including corticopontine (CPn) neurons, that project to deep subcortical targets, and intratelencephalic neurons, including transhemispheric commissural (COM) neurons, that primarily target structures within the telencephalon. Acting at M1-subtype muscarinic receptors, ACh preferentially boosts the excitability of CPn neurons relative to COM neurons (Dembrow et al., 2010; Baker et al., 2018), while 5-HT inhibits CPn neurons via G_i/o_-coupled 5-HT_1A_ (1A) receptors, but promotes action potential generation in COM neurons via Gq-coupled 5-HT_2A_ (2A) receptors (Avesar and Gulledge, 2012).

Corticoamygdalar (CAm) neurons in the prelimbic cortex that bilaterally target the basolateral amygdala (McDonald et al., 1996; Vertes, 2004; Gabbott et al., 2005) represent a less studied, but important, cortical projection that contributes to the expression of learned fear (Stevenson, 2011; Do Monte et al., 2016), and perhaps more generally to coping strategies in the face of external stressors (Varela et al., 2012). For instance, electrical stimulation of the prelimbic cortex promotes expression, and prevents extinction, of conditioned fear (Vidal-Gonzalez et al., 2006), while pharmacological inactivation of the prelimbic cortex impairs expression of conditioned freezing (Corcoran and Quirk, 2007; Sierra-Mercado et al., 2011; Almada et al., 2015). Indeed, unit activity (Burgos-Robles et al., 2009) and field potential oscillations (Dejean et al., 2016; Karalis et al., 2016) in the prelimbic cortex are correlated with animal freezing behavior in conditioned fear paradigms.

Accumulating evidence suggests that fear circuits may be regulated by 5-HT (for reviews, see Bauer, 2015; Bocchio et al., 2016). In rodents, fear conditioning is associated with 5-HT release in the medial prefrontal cortex (Yoshioka et al., 1995; Hashimoto et al., 1999), and acute administration of selective serotonergic reuptake inhibitors (SSRIs), which act to increase extracellular 5-HT levels, enhances the acquisition and expression of cued fear (Burghardt and Bauer, 2013; Ravinder et al., 2013). Studies targeting specific 5-HT receptor subtypes have found that activation of 1A receptors reduces, while 2A receptor stimulation enhances, conditioned fear responses. For instance, systemic injection of a 2A agonist (Zhang et al., 2013) enhances freezing in tests of cued and contextual fear memory in mice, while systemic activation of 1A receptors reduces freezing behavior in similar tests in rats (Inoue et al., 1996; Ohyama et al., 2016) or mice (Youn et al., 2009). Consistent with these findings, local activation of 1A receptors in the prelimbic cortex reduces conditioned fear responses and fear-potentiated startle (Ferreira and Nobre, 2014; Almada et al., 2015), whereas blockade of prelimbic 2A receptors impairs the expression of conditioned fear in a rat strain selectively bred to exhibit heightened anxiety (León et al., 2017). Together, these results suggest that serotonergic activation of 1A receptors in the prelimbic cortex may moderate fear responses, while engagement of 2A receptors may promote the expression of conditioned fear.

To better understand the physiology of CAm neurons, and the potential role of 5-HT in regulating cortical output to the amygdala, we used retrograde labeling to identify and record from CAm neurons in slices of mouse prelimbic cortex. Here we describe the physiological and morphological properties of these neurons, and their responsivity to 5-HT, in the context of other previously studied prelimbic projection neuron subtypes. Our results demonstrate that prelimbic CAm neurons are in many ways similar to, and overlap with, COM neurons, and suggest that 5-HT, acting on 2A receptors, may promote CAm output to the amygdala.

## Methods

### Animals

Experiments utilized tissue from 8-12-week-old male and female C57BL/6J mice (Jackson Laboratories, Bar Harbor, ME) according to methods approved by the Institutional Animal Care and Use Committee of Dartmouth College. Animals had access for food and water *ad libitum* and were housed on a 12:12 hour light:dark cycle.

### Retrograde Labeling

Red or green fluorescent beads (Retrobeads, Lumafluor Inc.) were injected unilaterally into the amygdala to label CAm projection neurons using age appropriate coordinates (Paxinos and Franklin, 2004). In some animals, an additional injection of retrobeads was made into the ipsilateral medial prefrontal cortex (mPFC) to double-label CAm/COM dual-projection neurons. Animals were anesthetized throughout surgeries with vaporized isoflurane (∼2%). Following craniotomy, a microsyringe was lowered into the brain region of interest, and 300 nL of undiluted Retrobead solution was injected over a 10 min period. In some experiments a nanoject (Drummond scientific) was used to administer the microbeads instead of a microsyringe. Animals were allowed to recover from surgery for at least 72 hours before use in electrophysiological experiments. Locations of dye injections were confirmed *post hoc* in coronal sections of the amygdala or mPFC.

### Slice Preparation

Following isoflurane anesthesia and decapitation, brains were quickly removed into artificial cerebral spinal fluid (aCSF) containing, in mM: 125 NaCl, 25 NaHCO_3_, 3 KCl, 1.25 NaH_2_PO_4_, 0.5 CaCl_2_, 5 MgCl_2_, and 25 glucose, saturated with 95% O_2_ / 5% CO_2_. Coronal brain slices (250 µm thick) of the mPFC or amygdala were cut using a Leica VT 1200 slicer and placed in a holding chamber filled with ACSF containing 2 mM CaCl_2_ and 1 mM MgCl_2_. Slices were stored at 35°C for ∼45 min, then kept at room temperature for up to 8 hours before use in experiments.

### Electrophysiology

Slices were placed in a recording chamber on a fixed-stage Olympus BX51WI microscope and continuously perfused with oxygenated ACSF heated to 35–36°C. Epifluorescence illumination (470 or 530 nm LEDs) was used to identify CAm or CAm/COM neurons in the mPFC, and whole-cell current-clamp recordings were made from bead-labeled neurons using patch pipettes (5 – 7 MΩ) filled with, in mM: 135 K-gluconate, 2 NaCl, 2 MgCl_2_, 10 HEPES, 3 Na_2_ATP, and 0.3 NaGTP (pH 7.2 with KOH). Data were acquired with Axograph software (Axograph Company) using BVC-700 amplifiers (Dagan Corporation) and ITC-18 digitizers (HEKA Instruments). Membrane potentials were sampled at 25 kHz, filtered at 10 kHz, and corrected for a liquid junction potential of +12 mV. Hyperpolarization-activated cyclic-nucleotide gated (HCN)-channel-mediated rebound (i.e., “sag” potentials) were measured as the percent difference in the peak hyperpolarization and the steady state voltage (both relative to resting membrane potentials) in response to negative current injection set to hyperpolarize the neuron by approximately 20 mV.

5-HT (100 µM) was dissolved in ACSF and loaded into patch pipettes placed ∼50 µm from targeted somata. Neurons were classified as 5-HT-inhibited, -excited, -biphasic, or -non-responsive based on their response to focal application of 5-HT, which was delivered for 1 s at ∼15 PSI during periods of action potential (AP) generation (~5 Hz) evoked by somatic DC current injection. Serotonergic inhibition was quantified as the duration of cessation of action potential generation, while excitatory responses were quantified as the percent increase in the mean instantaneous spike frequency (ISF; the inverse of the time interval between each AP) measured over the 500 ms following the peak post-5-HT ISF, relative to the mean pre-5-HT ISF baseline (10 s). Biphasic 5-HT responses were defined as responses having both brief inhibition (lasting at least 10 times the average baseline interspike interval) and a subsequent increase in ISF of at least 1 Hz above baseline levels. Mean “population” responses to 5-HT during DC-current-induced action potential firing were made by resampling individual ISF plots at 2 Hz and then averaging those ISF plots across tested neurons. In some experiments, 5-HT receptors were selectively blocked with 1A (WAY 100635, 30 nM; Sigma–Aldrich) and/or 2A (MDL 11939, 500 nM; Tocris) receptor antagonists.

In some experiments, somatic current injection were used to simulate excitatory synaptic input, as previously described (Stephens et al., 2014). Because of intrinsic cell-to-cell variability in input resistance and excitability, for each neuron the synaptic waveform was scaled in amplitude to evoke ∼8 action potentials during baseline trials. The simulated synaptic current was then delivered 29 times at 3 s intervals, the exception being the sixth trial which was delayed 3 s due to application of 5-HT (100 µM, 1 s).

### Morphological Analyses and Single-cell Imaging

In some experiments, biocytin (7 mg/ml; Tocris) was included in the patch-pipette solution. Following recordings, slices were placed in a phosphate buffered saline (PBS) solution containing 4% paraformaldehyde for 24 hours. Slices were then rinsed three times for fifteen minutes in PBS and placed in a PBS solution containing 0.25% Triton X-100 and avidin conjugated to either Alexa Fluor®-594 or Alexa Fluor®-488 for 24 hours (20 µg/ml; Invitrogen). Slices were then rinsed with PBS three times for fifteen minutes each, dried, and mounted on slides in FluorSave (EMD Chemicals). Cells were imaged using a 2-photon microscope (Prairie Technologies) or a Zeiss LSM 510 confocal microscope (Carl Zeiss). Data from COM and CPn neurons used for comparisons were previously published in Avesar and Gulledge (2012) and Stephens et al. (2014). Morphological measurements included somatic distance from the pia, maximum horizontal width of the apical tuft, the number of dendritic branch points in the tuft, and the number of primary and oblique dendrites. Tracings of neurons were made using NeuronJ from z-stack projections.

### Statistical Analyses

Unless otherwise noted, all data are presented as mean ± standard deviation. Comparisons across cell groups utilized one-way ANOVAs with Šidák post-tests, while comparisons within groups was accomplished using 2-tailed Student’s *t*-tests (paired or unpaired), or repeated measures ANOVA, as appropriate. Comparisons of response proportions between groups utilized a Fisher’s exact test. Significance was defined as *p* < 0.05.

## Results

### Physiological and morphological characteristics of CAm neurons

Retrograde-labeled pyramidal neurons projecting to either the ipsilateral (iCAm neurons; n = 24) or contralateral (cCAm neurons; n = 24) amygdala were targeted for whole-cell recordings in layer 5 of the mouse prelimbic cortex. For each neuron, measurements were made of resting membrane potential (RMP), input resistance (R_N_), and HCN-channel-mediated rebound “sag” potentials (% sag; see Methods; **Table 1** and **Figure 1**). CAm neuron physiology was not sex-dependent (**Table 1**), and RMP and HCN responses were not dependent on the laterality of amygdalar projection (**Table 1**). However, ipsilaterally projecting CAm neurons had a significantly higher input resistance than contralateral CAm neurons (p = 0.031; Student’s *t*-test; **Table 1**). The physiological properties of CAm neurons, as a whole, were somewhat distinct from those previously observed in layer 5 COM and CPn neurons (Avesar and Gulledge, 2012; Stephens et al., 2014). While CAm neurons resembled COM neurons in having larger R_N_s (at 172 ± 54 MΩ; n = 48; *p* < 0.001; Student’s *t*-test) and smaller HCN-dependent sag responses (at 9.8 ± 4.8%; *p* < 0.001) than CPn neurons (n = 58; **Table 1**), the RMPs of CAm neurons (−78 ± 5 mV) were similar to those in CPn neurons (−77 ± 3 mV; p = 0.73), and more depolarized than those in COM neurons (−80 ± 5 mV; n = 64; p = 0.039), likely reflecting their enhanced sag potentials relative to those of COM neurons (6.4 ± 4.6%; *p* < 0.001; see **Table 1** and **Figure 1C**).

**Table 1:**
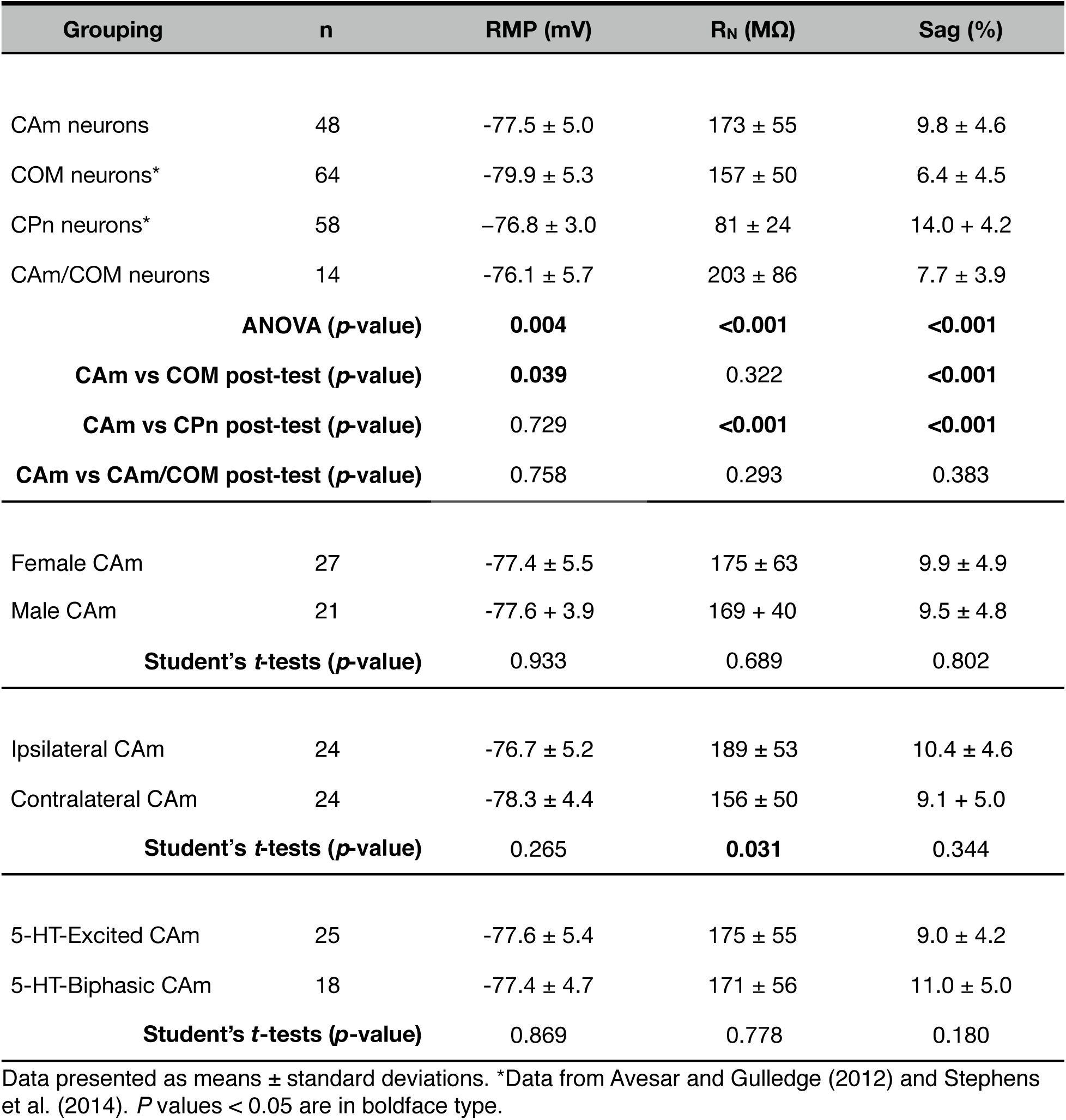
Physiological properties of layer 5 neurons

| Grouping | n | RMP (mV) | R<sub>N</sub> (MΩ) | Sag (%) |
| --- | --- | --- | --- | --- |
| CAM neurons | 48 | -77.5 ± 5.0 | 173 ± 55 | 9.8 ± 4.6 |
| COM neurons* | 64 | -79.9 ± 5.3 | 157 ± 50 | 6.4 ± 4.5 |
| CPn neurons* | 58 | -76.8 ± 3.0 | 81 ± 24 | 14.0 ± 4.2 |
| CAM/COM neurons | 14 | -76.1 ± 5.7 | 203 ± 86 | 7.7 ± 3.9 |
| **ANOVA (*p*-value)** |  | **0.004** | **<0.001** | **<0.001** |
| **CAM vs COM post-test (*p*-value)** |  | **0.039** | 0.322 | **<0.001** |
| **CAM vs CPn post-test (*p*-value)** |  | 0.729 | **<0.001** | **<0.001** |
| **CAM vs CAM/COM post-test (*p*-value)** |  | 0.758 | 0.293 | 0.383 |
| Female CAM | 27 | -77.4 ± 5.5 | 175 ± 63 | 9.9 ± 4.9 |
| Male CAM | 21 | -77.6 ± 3.9 | 169 ± 40 | 9.5 ± 4.8 |
| **Student's *t*-tests (*p*-value)** |  | 0.933 | 0.689 | 0.802 |
| Ipsilateral CAM | 24 | -76.7 ± 5.2 | 189 ± 53 | 10.4 ± 4.6 |
| Contralateral CAM | 24 | -78.3 ± 4.4 | 156 ± 50 | 9.1 ± 5.0 |
| **Student's *t*-tests (*p*-value)** |  | 0.265 | **0.031** | 0.344 |
| 5-HT-Excited CAM | 25 | -77.6 ± 5.4 | 175 ± 55 | 9.0 ± 4.2 |
| 5-HT-Biphasic CAM | 18 | -77.4 ± 4.7 | 171 ± 56 | 11.0 ± 5.0 |
| **Student's *t*-tests (*p*-value)** |  | 0.869 | 0.778 | 0.180 |

Data presented as means ± standard deviations. \*Data from Avesar and Gullledge (2012) and Stephens et al. (2014). *P* values < 0.05 are in boldface type.

**Figure 1.**
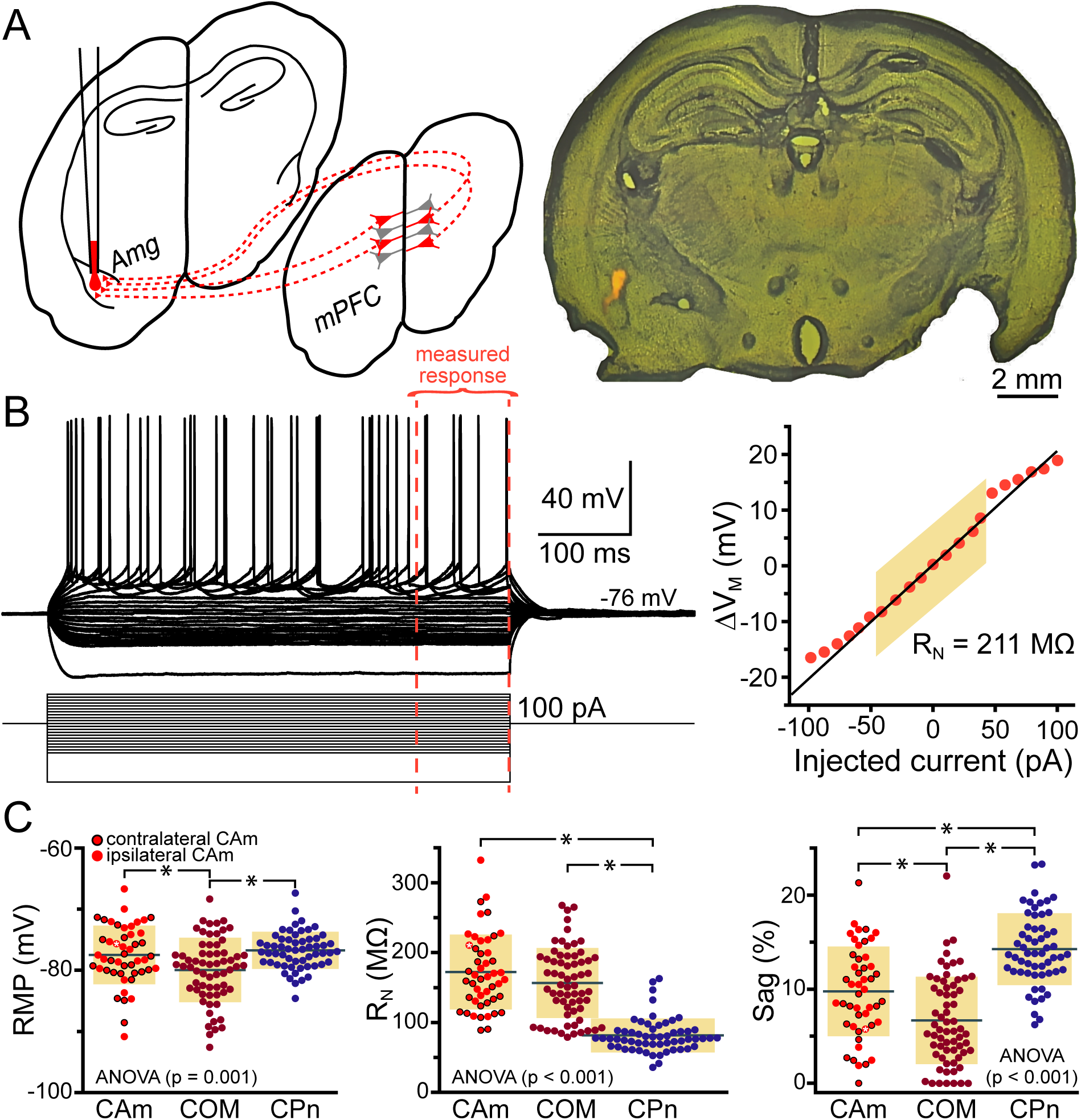
Physiological properties of corticoamygdalar neurons. **A**. *Left* - Schematic of bilateral retrograde labeling of corticoamygdalar (CAm) neurons in the medial prefrontal cortex. *Right* - a coronal section of a mouse brain showing the amygdala containing fluorescent retrograde tracers. **B**. Voltage traces (*top left*) of a L5 CAm neuron in response to a family of positive and negative current injections (*bottom left*). Red dashed lines show the final 100 ms of the voltage responses. At right is a plot of the voltage-current relationship as measured across the final 100 ms of the voltage responses. The shaded area shows the linear region of the relationship that was used to calculate input resistance (R_N_). **C.** Plots of individual measurements of resting membrane potential (RMP; *left*), R_N_ (*middle*), and HCN-channel dependent voltage sag (*right*) for CAm (n = 48), COM (n = 64), and CPn (n = 58) neurons. Black horizontal lines indicate means, shaded regions indicate ± 1 standard deviation. Symbols with black borders indicate contralateral CAm neurons. Symbols with white asterisks indicate the data from the cell shown in **B**. Data for COM and CPn neurons aggregated from Avesar and Gulledge (2012) and Stephens et al., (2014). Black asterisks indicate *p* < 0.05, Student’s *t*-test.

Morphologically, layer 5 CAm neurons (n = 21) resembled COM, rather than CPn, neurons (**Figure 2**). Both CAm and COM neurons had fewer primary (mean of 7.1 ± 1.5; *p* = 0.020, Šidák post-test) and oblique dendrites (7.4 ± 4.1; *p* = 0.001) than CPn neurons (n = 10; **Table 2** and **Figure 2B**). While cells of all projection subtypes exhibited similar apical tuft widths (e.g., 289 ± 126 µm for CAm neurons; *p* = 0.14, ANOVA), CAm and COM (n = 10) neuron somata tended to be slightly more superficial (at 339 ± 18 µm and 332 ± 18 um from the pia; *p* = 0.029 and 0.010; Šidák post-tests for CAm and COM neurons, respectively, vs CPn neurons) than the somata of CPn neurons (405 ± 13 µm; **Figure 2B** and **Table 2**). Thus, while CAm neurons were in some ways distinct from COM and CPn neurons (e.g, in their HCN-mediated responses), their overall physiological and morphological characteristics were more similar to COM neurons.

**Table 2:** Comparison of morphological properties of layer 5 projection neurons

| Grouping | n | Depth from pia ($\mu\text{m}$) | # primary dendrites | # oblique dendrites | Apical tuft width ($\mu\text{m}$) |
| --- | --- | --- | --- | --- | --- |
| CAM neurons | 21 | 335 ± 78<sup>†</sup> | 7.1 ± 1.5 | 7.4 ± 4.1 | 289 ± 126 |
| COM neurons* | 10 | 332 ± 58 | 6.7 ± 2.4 | 4.6 ± 2.3 | 211 ± 95 |
| CPn neurons* | 10 | 405 ± 41 | 9.2 ± 2.8 | 13.3 ± 3.5 | 302 ± 96 |
| **ANOVA (*p*-value)** |  | **0.026** | **0.018** | **< 0.001** | 0.140 |
| **CAM vs COM post-test (*p*-value)** |  | 0.999 | 0.922 | 0.165 |  |
| **CAM vs CPn post-test (*p*-value)** |  | **0.043** | **0.030** | **0.002** |  |

Data presented as means ± standard deviations. \*Data from Avesar and Gullledge (2012). <sup>†</sup>n = 19 for depth from pia. *P* values < 0.05 are in boldface type.

**Figure 2.**
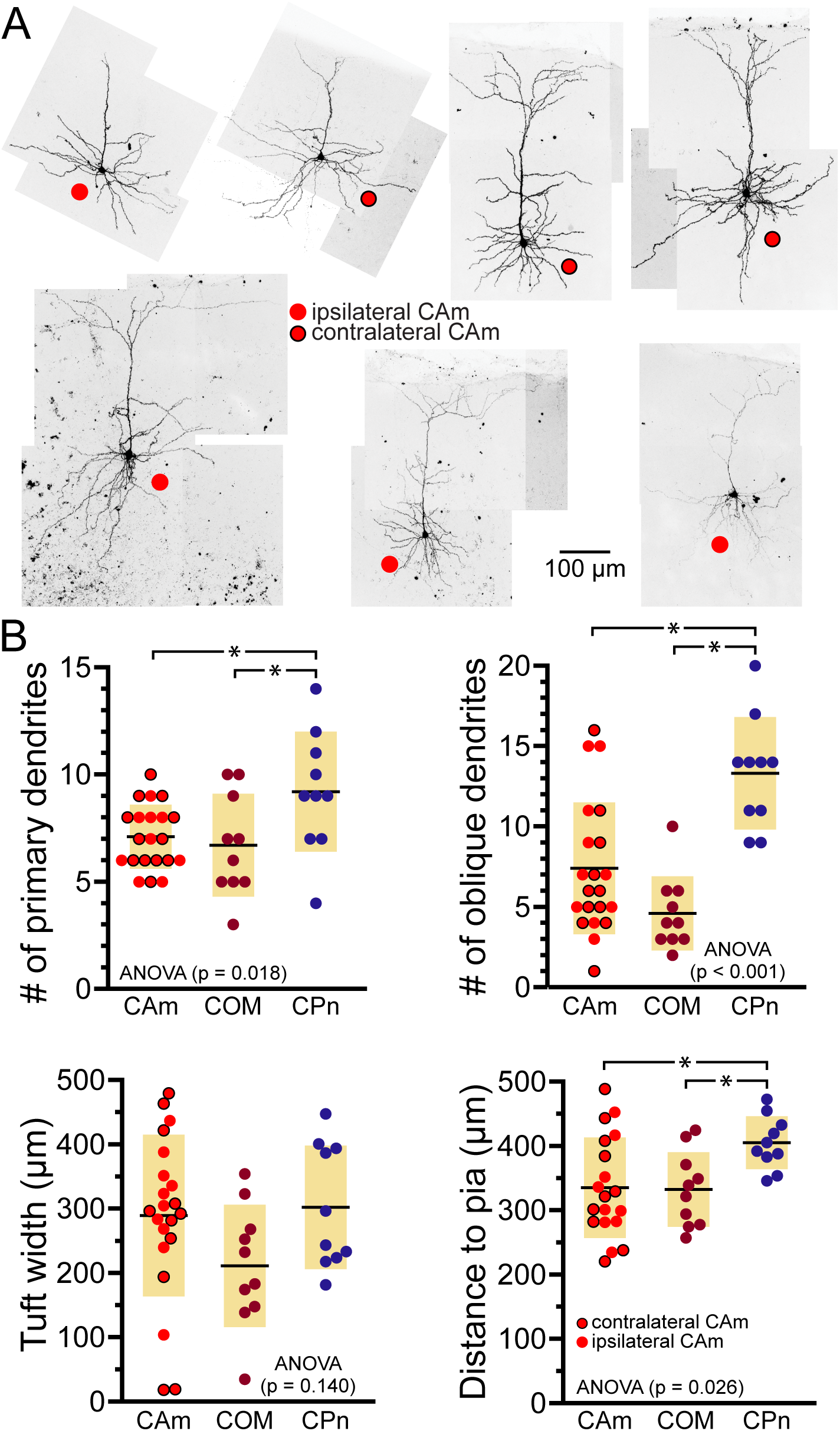
Morphological characteristics of CAm neurons. **A.** Images of CAm neurons filled with biocytin and conjugated to Alexa-488. Red circles without borders indicate CAm neurons projecting to the ipsilateral amygdala, while those with black borders indicate contralaterally projecting CAm neurons. **B.** Plots of morphological features for ipsilaterally and contralaterally projecting CAm neurons (n = 21), COM (n = 10), and CPn (n = 10) neurons. Black horizontal lines indicate means, shaded regions indicate ± 1 standard deviation. Morphological measurements for COM and CPn neurons were previously published in Avesar and Gulledge (2012). Asterisks indicate *p* < 0.05, Student’s *t*-test.

### CAm neurons are excited by 5-HT via 2A receptors

CAm neurons also resembled COM neurons in their responses to focal application of 5-HT during periods of current-induced action potential generation (**Figure 3; Table 3**). 5-HT enhanced the frequency of action potential generation in 25 of 48 CAm neurons (52%), and generated biphasic responses in 18 CAm neurons (38%; **Figure 3A**, **B**). Four CAm neurons were non-responsive to 5-HT, and one CAm neuron exhibited a purely inhibitory response to 5-HT. The proportions of CAm excited and biphasic neurons were statistically similar in 24 iCAm and 24 cCAm neurons (*p* = 0.36; Fisher’s exact test; **Figure 3C**), suggesting that 5-HT influences CAm neurons independently of the laterality of their amygdalar projections. Indeed, the magnitudes of serotonergic excitation were similar between iCAm (n = 11) and cCAm (n = 14) 5-HT-excited neurons (mean excitations of 84 ± 63 and 121 ± 115%, respectively; *p* = 0.32; Student’s *t*-test). Excitatory responses in iCAm (n = 11) and cCAm (n = 7) 5-HT-biphasic neurons (mean excitations of 120 ± 131 and 85 ± 49%, respectively; **Figure 3D**) were also similar to each other (*p* = 0.44) and to the magnitude of excitation observed across all 5-HT-excited CAm neurons (98 ± 71%; n = 25; *p* = 0.77, ANOVA). Further, serotonergic biphasic inhibitions in iCAm and cCAm neurons were of similar durations (at 16 ± 8 and 19 ± 5 ms, respectively; *p* = 0.64; Student’s *t*-test; **Figure 3E**). These results suggest that 5-HT responses in CAm neurons are not dependent on projection laterality. Additionally, 5-HT responses in CAm neurons were not sex-dependent (**Table 3**).

**Table 3:** Serotonergic responses in CAm neurons

| Grouping | n | 5-HT response | Excitation (% increase in ISF) | Duration of inhibition (s) |
| --- | --- | --- | --- | --- |
| L5 COM neurons* | 43 | Excited | 142 ± 101 | NA |
|  | 21 | Biphasic | 153 ± 78 | 17 ± 9 |
| L5 CAm neurons | 25 | Excited | 98 ± 97 | NA |
|  | 18 | Biphasic | 111 ± 102 | 18 ± 11 |
| L2/3 CAm neurons | 7 | Excited | 93 ± 71 | NA |
|  | 6 | Biphasic | 148 ± 101 | 27 ± 19 |
| L5 CAm/COM neurons | 9 | Excited | 54 ± 34 | NA |
|  | 4 | Biphasic | 70 ± 45 | 16 ± 7 |
| **ANOVA** |  |  | 0.077 | 0.249 |
| Ipsilateral L5 CAm | 11 | Excited | 84 ± 63 | NA |
| Contralateral L5 CAm | 14 | Excited | 121 ± 115 | NA |
| **Student's *t*-test (*p*-value)** |  |  | 0.321 |  |
| Ipsilateral L5 CAm | 11 | Biphasic | 120 ± 131 | 16 ± 8 |
| Contralateral L5 CAm | 7 | Biphasic | 85 ± 49 | 19 ± 15 |
| **Student's *t*-tests (*p*-value)** |  |  | 0.436 | 0.643 |
| Female L5 CAm | 15 | Excited | 108 ± 83 | NA |
| Male L5 CAm | 10 | Excited | 99 ± 116 | NA |
| **Student's *t*-test (*p*-value)** |  |  | 0.837 |  |
| Female L5 CAm | 9 | Biphasic | 120 ± 142 | 20 ± 15 |
| Male L5 CAm | 9 | Biphasic | 93 ± 58 | 15 ± 5 |
| **Student's *t*-tests (*p*-value)** |  |  | 0.611 | 0.315 |

Data presented as means ± standard deviations. \*Data from Avesar and Gullledge (2012) and Stephens et al. (2014). NA = not applicable.

**Figure 3.**
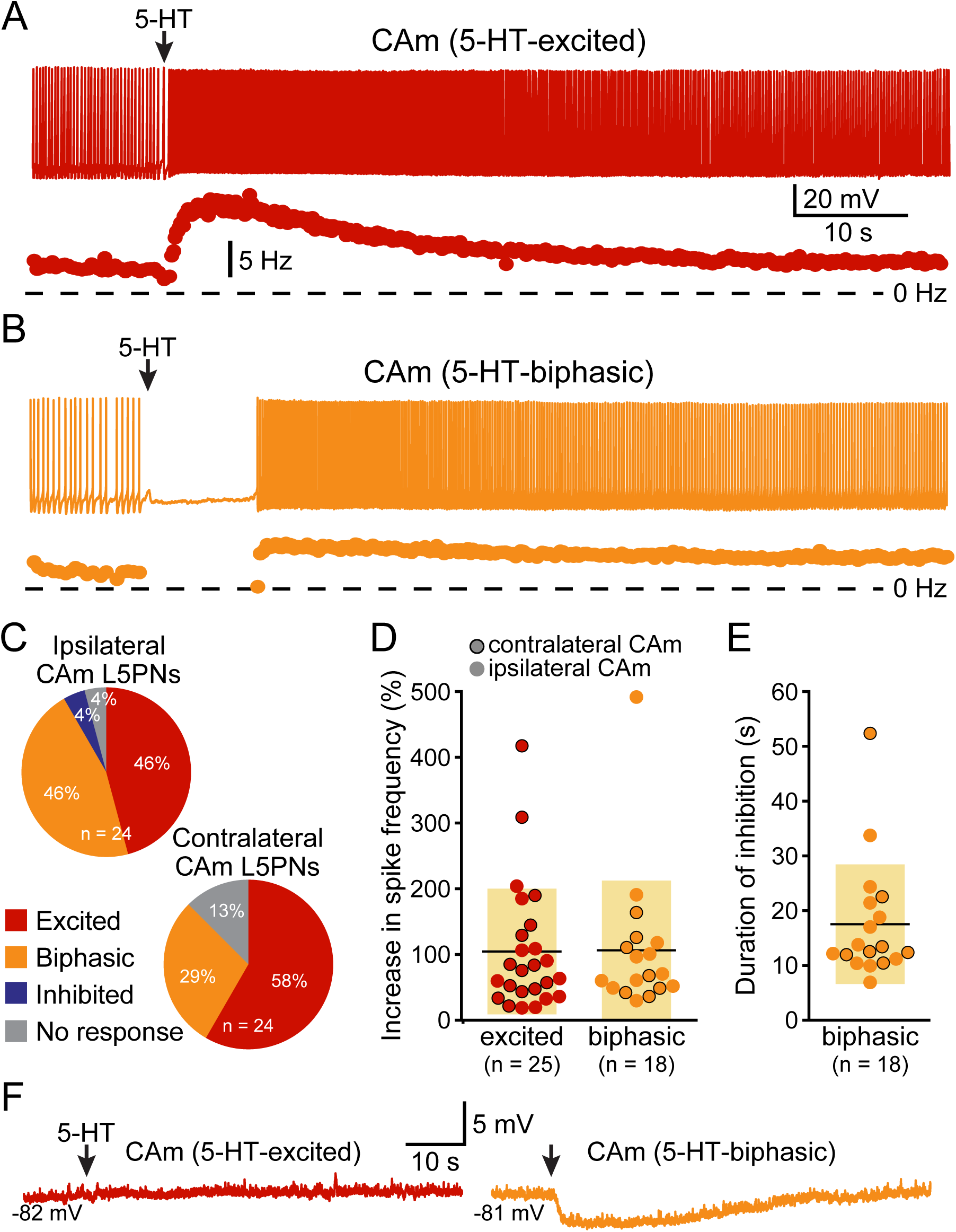
5-HT excites CAm neurons. **A.** Voltage trace (*top*) and plot of instantaneous spike frequency (ISF; *bottom*) of action potentials in a CAm neuron during prolonged somatic DC current injection. Focal application of 100 µM 5-HT (1 s; arrow) enhanced action potential frequency in this neuron. Dashed line indicates zero Hz. **B.** Voltage trace (*top*) and plot of ISF (*bottom*) over time in a CAm neuron exhibiting a biphasic (inhibition followed by excitation) response to focal application of 5-HT (arrow). **C.** Graphs showing the proportions of ipsilaterally and contralaterally projecting layer 5 CAm neurons that exhibited excitatory (red), biphasic (yellow), inhibitory (blue) responses, or no response at all (gray). **D.** Plots of the magnitudes of excitatory responses to 5-HT (% relative to baseline firing rates) for ipsilaterally (non-bordered symbols) and contralaterally (black-bordered symbols) projecting 5-HT-excited (red) and 5-HT-biphasic (yellow) CAm neurons. **E.** Plots of the duration of 5-HT-induced spike-cessation in ipsilaterally and contralaterally projecting 5-HT-biphasic CAm neurons. **F.** Voltage traces of a 5-HT-excited CAm neuron (red, left) and a 5-HT-biphasic neuron (yellow, right) during 5-HT application at resting membrane potentials (−82 and −81 mV, respectively).

5-HT responses in layer 5 CAm neurons were qualitatively and quantitatively similar (in both proportions and magnitudes) to 5-HT responses previously reported in COM neurons (see **Table 3**). Further, as previously reported in COM neurons (Stephens et al., 2014), serotonergic excitation of CAm neurons required additional depolarizing drive, as 5-HT failed to depolarize 5-HT-excited CAm (n = 8) or 5-HT-biphasic (n = 7) neurons from RMPs (**Figure 3F**). In 5-HT-excited CAm neurons, the mean change in membrane potential relative to RMPs was +0.1 ± 0.5 mV (*p* = 0.54, paired Student’s *t*-test). In 5-HT-biphasic CAm neurons, focal application of 5-HT at RMPs led to transient hyperpolarizations (of 3.5 ± 0.9 mV; *p* < 0.001) that lasted 27 ± 11 s, but no subsequent depolarizations (mean peak membrane potential relative to RMPs was +0.4 ± 1.1 mV; *p* = 0.43). Thus, serotonergic stimulation, on its own, was not sufficient to excite CAm neurons.

Serotonergic excitation of COM neurons is mediated by 2A receptors, whereas biphasic responses involve both 1A-mediated inhibition and 2A-driven excitation (Avesar and Gulledge, 2012; Stephens et al. 2014). To test whether 2A receptors also mediate serotonergic excitation of CAm neurons, we measured 5-HT responses in 5-HT-excited and 5-HT-biphasic CAm neurons before and after bath application of the selective 2A receptor antagonist MDL 11939 (500 nM). In both 5-HT-excited (n = 8) and 5-HT-biphasic (n = 7) CAm neurons, MDL blocked serotonergic excitation (**Figure 4**). In a subset of 5-HT-biphasic neurons (n = 3), the inhibitory portion of biphasic responses was blocked by WAY 100635 (30 nM), a 1A antagonist (**Figure 4C**). These results confirm that serotonergic responses in prelimbic CAm neurons are mediated by the same receptor subtypes that gate serotonergic excitation and inhibition in COM and CPn neurons (Avesar and Gulledge, 2012; Stephens et al. 2014).

**Figure 4.**
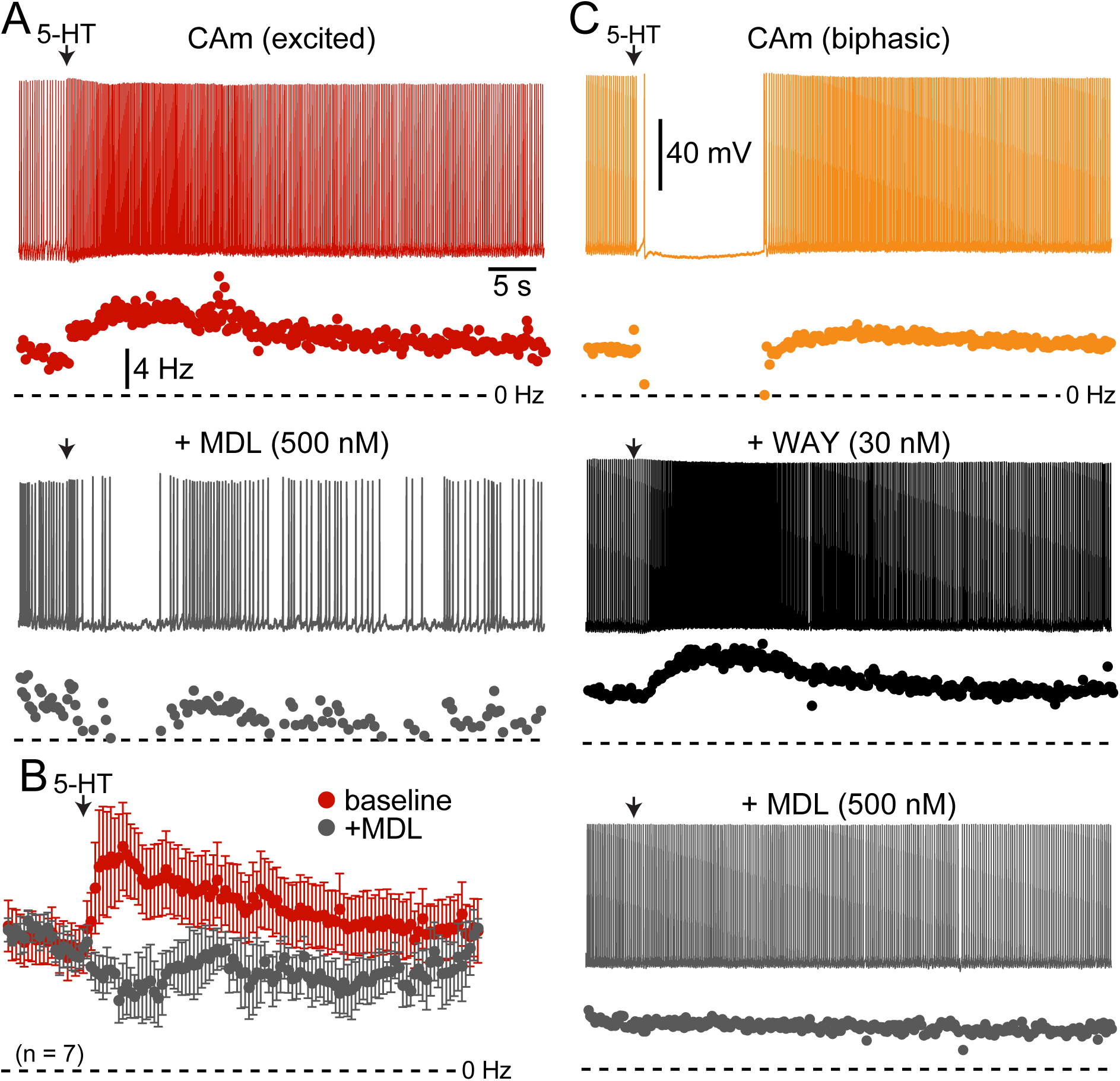
Serotonergic responses in CAm neurons are mediated by 1A and 2A receptors. **A.** Excitatory serotonergic response of a layer 5 CAm neuron (red voltage trace and plot of instantaneous spike frequency [ISF], *top*) that was subsequently blocked by bath application of the 2A antagonist MDL 11939 (500 nM; gray voltage trace and ISF plot). Arrows indicate timing of 5-HT application. **B.** Population response (mean ISF ± SEM) to 5-HT for seven 5-HT-excitd CAm neurons before (red) and after (gray) addition of 500 nM of MDL 11939. **C.** Voltage traces (top) and ISF plots (bottom) showing responses to 5-HT (arrow) in a biphasic layer 5 CAm neuron under baseline conditions (yellow trace/plot) and after sequential addition of the 1A receptor antagonist WAY 100635 (30 nM; black trace/plot) and the 2A antagonist MDL 11939 (500 nM; gray trace/plot).

### 5-HT enhances CAm neuron output to simulated synaptic input

To test whether 5-HT can functionally enhance the output of CAm neurons driven by a more physiological stimulus, we repeatedly delivered a barrage of simulated synaptic current (see Methods) over 29 trials (3 s inter-trial intervals), and measured the number of action potentials generated in each trial, as well as the RMP just prior to the barrage (**Figure 5**). Following the initial five baseline trials, 5-HT was applied for 1 s before synaptic barrages were resumed for 24 additional trials. In 5-HT-excited CAm neurons (n = 14), simulated synaptic barrages generated a mean of 7.6 ± 1.2 action potentials during the five baseline trials. Following 5-HT application, there was an increase of 2.4 ± 1.9 action potentials per trial, bringing the mean response to 10.0 ± 2.1 action potentials across the subsequent five trials (*p* < 0.001; repeated measures ANOVA; **Figure 5B**, red symbols). Conversely, 5-HT had little impact on action potential output in biphasic CAm neurons (mean change of −1.6 ± 4 action potentials, relative to their baseline response of 8.9 ± 1.7 action potentials; *p* = 0.31, **Figure 5B**, yellow symbols). Since 5-HT generated a small, but significant (*p* = 0.024), depolarization of the RMP in 5-HT-excited CAm neurons (peaking with a mean of 1.2 ± 1.3 mV on trial 11, about 6 s after 5-HT application, and only after application of suprathreshold current input), in a subset of 5-HT-excited neurons (n = 8) we tested whether depolarization alone could account for the additional action potentials produced after 5-HT application. In these control experiments, in which 7.1 ± 1.5 spikes were generated in baseline trials, DC current was used to continuously depolarize neurons by 1.8 ± 0.7 mV (*p* < 0.001) during the remaining 24 trials. This depolarization alone failed to significantly increase the number of action potentials generated by the simulated synaptic input (mean of 7.7 ± 2.2 action potentials; *p* = 0.12). Together, these results confirm previous findings in COM neurons (Stephens et al., 2014), and suggest that 5-HT will preferentially enhance the output of 5-HT-excited CAm neurons that are receiving coincident suprathreshold synaptic drive.

**Figure 5.**
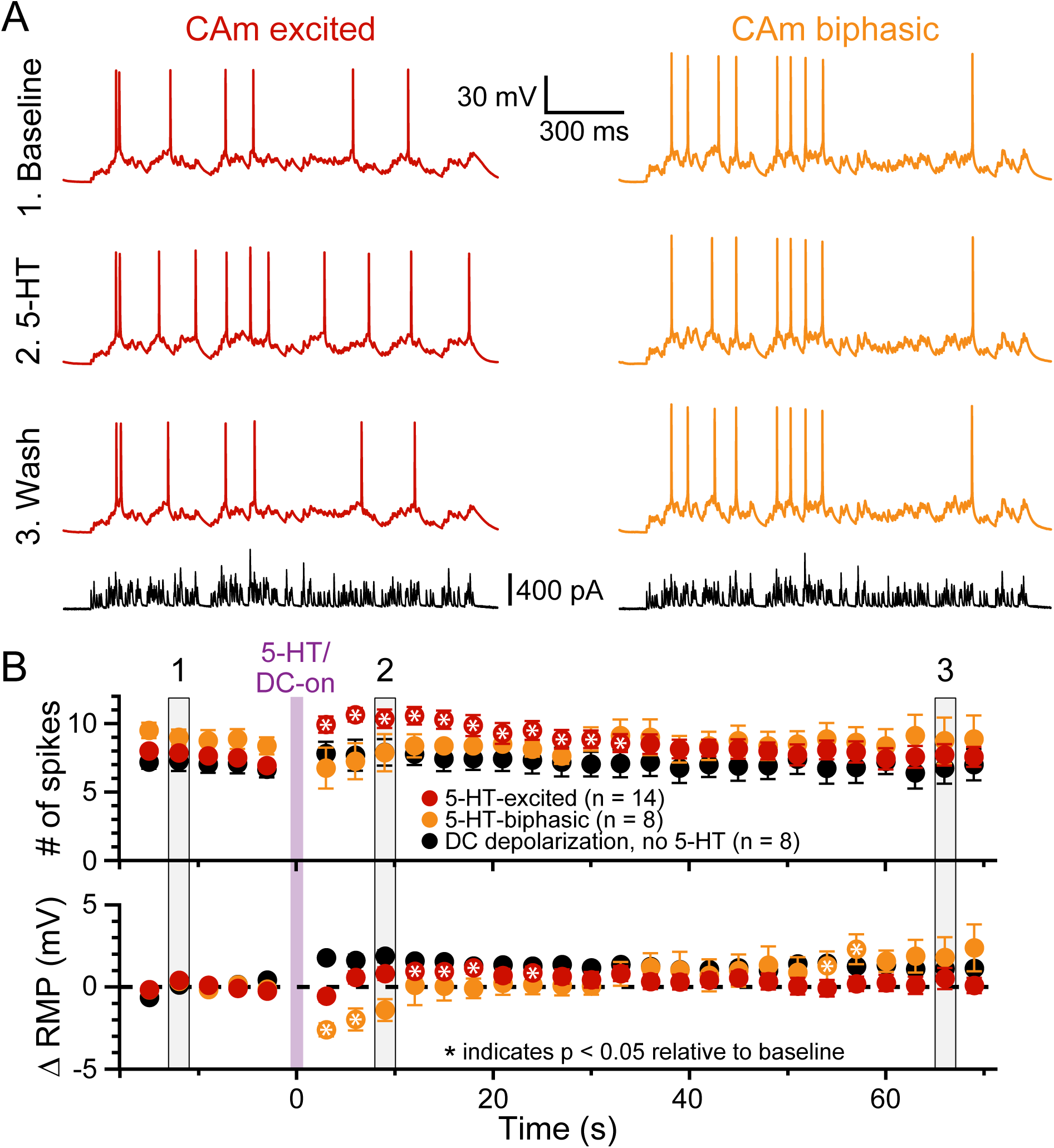
5-HT enhances CAm neuron output in response to simulated synaptic input. **A.** Voltage traces from 5-HT-excited (red traces) and 5-HT-biphasic (yellow traces) CAm neurons in response to a simulated barrage of synaptic input (black traces) delivered in (1) baseline conditions (*top* traces), (2) after focal application of 5-HT (middle traces), and (3) about 1 minute later (lower voltage traces). **B.** Population responses (means ± SEMs) for 5-HT-excited (red symbols; n = 14) and 5-HT-biphasic CAm neurons (yellow symbols; n = 8). Black symbols show data from a subset of 5-HT-excited neurons (n = 8) receiving DC current, instead of 5-HT, to depolarize the cells 1-2 mV following 5 baseline applications of simulated synaptic current. The mean number of action potentials in response to the simulated synaptic barrage are plotted over time in the top graph, while the lower graph plots the mean changes in resting membrane potentials (RMPs), relative to baseline values, as measured for each trial immediately before simulated synaptic current application. Asterisks indicates *p* < 0.05 relative to baseline measurements from repeated measures ANOVA. Grey vertical bars (1, 2, and 3) indicate the trials represented in **A**.

### 5-HT generates biphasic and excitatory responses in L2/3 CAm neurons

Although our study focuses on layer 5 pyramidal neurons, CAm neurons are present across cortical lamina (Little and Carter, 2013; Song et al., 2015; Cheriyan et al., 2016). To determine whether 5-HT regulates the excitability of superficial CAm neurons, we recorded from Retrobead-labeled CAm neurons in layer 2/3 of the prelimbic cortex (n = 13). Layer 2/3 CAm neurons had physiological properties comparable to those in layer 5, with mean RMPs of −77 ± 7 mV (*p* = 0.98 vs 48 layer 5 CAm; Student’s *t*-test), input resistances of 208 ± 82 MΩ (*p* = 0.61 vs layer 5 CAm), and sag potentials of 7.3 ± 3.8% (*p* = 0.30 vs layer 5 CAm). Layer 2/3 CAm neurons exhibited excitatory (n = 7) or biphasic (n = 6) responses to focally applied 5-HT (**Figure 6**; **Table 3**) that occurred in proportions (*p* = 1.00; Fisher’s exact test) and magnitudes (mean increases above baseline ISF of 93 ± 71% and 148 ± 101% for 5-HT-excited and 5-HT-biphasic L2/3 CAm neurons, respectively; *p* = 0.22 and 0.91, when compared to layer 5 CAm neurons) consistent with those observed in layer 5 CAm and COM neurons. Thus, serotonergic regulation of CAm neuron excitability appears to be conserved across cortical layers in the prelimbic cortex.

**Figure 6.**
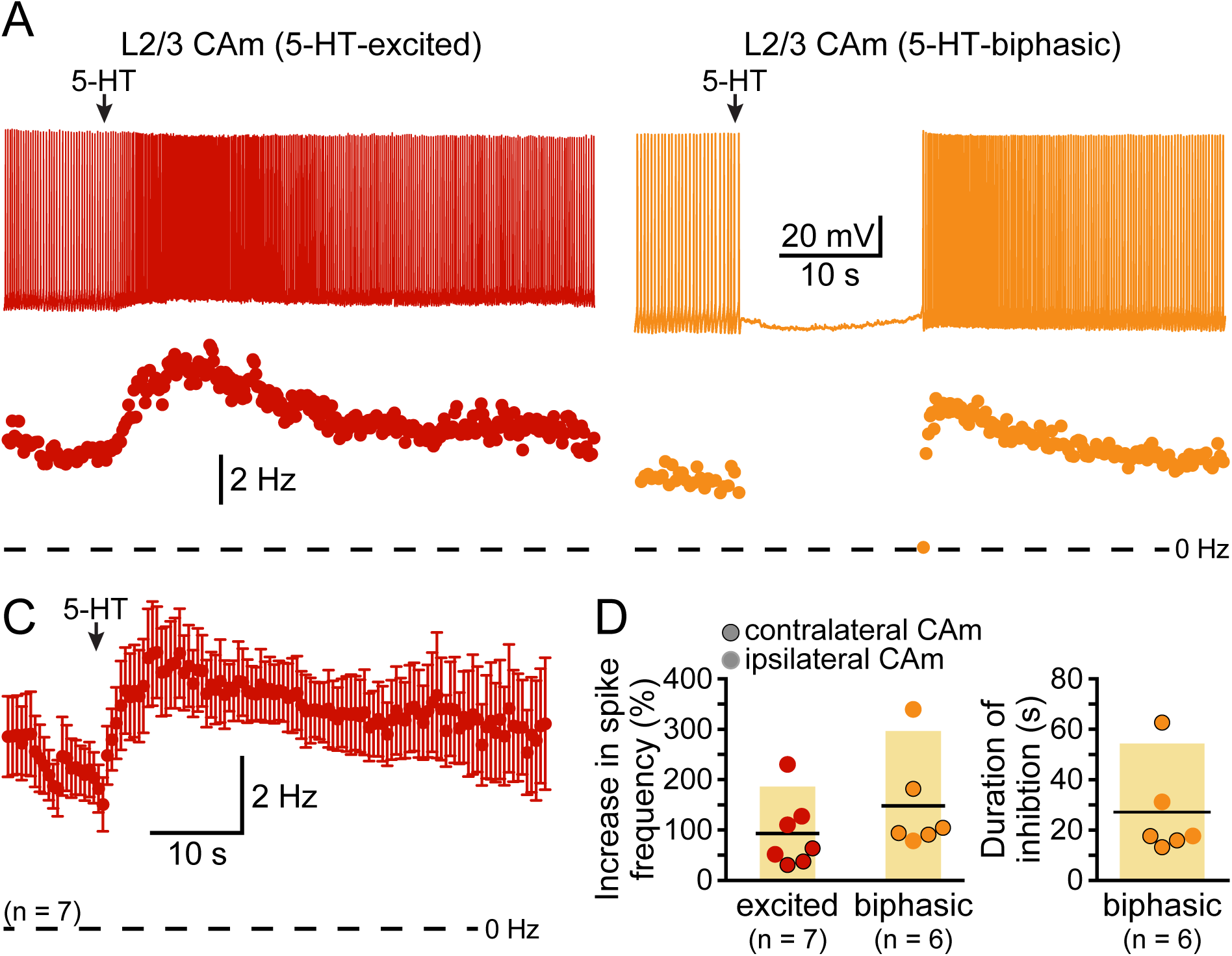
Serotonergic regulation of layer 2/3 CAm neurons. **A.** Voltage traces (*top*) and plots of instantaneous spike frequencies (ISFs; *bottom*) of action potentials in two CAm neurons in layer 2/3 of the prelimbic cortex experiencing continuous somatic DC current injection. Focal application of 100 µM 5-HT (1 s; arrows) increased the frequency of action potential genesis in one neuron (*left*), and a biphasic response in the other neuron (*right*). **B.** Population response (mean ISF ± SEM) to 5-HT for seven 5-HT-excited layer 2/3 CAm neurons. **C.** Plots of the increase in spike frequency in response to 5-HT (% relative to baseline firing rates) for ipsilaterally (non-bordered symbols) and contralaterally (black-bordered symbols) projecting 5-HT-excited (red) and 5-HT-biphasic (yellow) layer 2/3 CAm neurons (*left*), and of the duration of 5-HT-induced spike-cessation in 5-HT-biphasic layer 2/3 CAm neurons (*right*).

### CAm neurons overlap with COM neurons

Based on their morphological and physiological similarities, we hypothesized that CAm and COM neurons may represent overlapping populations of layer 5 neurons in the mPFC. We tested this hypothesis by injecting retrobeads unilaterally into the contralateral amygdala and the mPFC to label CAm/COM dual projection neurons (**Figure 7A**). While CAm/COM neurons were much less numerous than single-labeled COM and CAm neurons, we recorded from 14 dual-labeled neurons, finding that the majority (64%, n = 9/14) were excited by 5-HT, with an additional 29% (n = 4/14) exhibiting biphasic responses to 5-HT. One CAm/COM double-labeled neuron (7%) exhibited a purely inhibitory response to 5-HT (**Figure 7B**, **C**). The magnitudes of excitatory responses in CAm/COM 5-HT-excited (54 ± 34%) and 5-HT-biphasic (70 ± 45%) neurons (**Figure 7D** and **Table 3**) were statistically similar to those observed in single-labeled 5-HT-excited CAm neurons (*p* = 0.13 and 0.51, for CAm/COM 5-HT-excited and 5-HT-biphasic neurons, respectively; Student’s *t*-tests). Similarly, the durations of inhibitory responses in CAm/ COM 5-HT-biphasic neurons (19 ± 5 s) were similar to those in single-labeled CAm neurons (*p* = 0.81). The physiological properties of CAm/COM neurons were also similar to those observed in single-labeled CAm neurons (**Table 1**). These data demonstrate that, in the mouse prelimbic cortex, CAm and COM neurons are overlapping neuron populations that share 2A-mediated serotonergic excitation.

**Figure 7.**
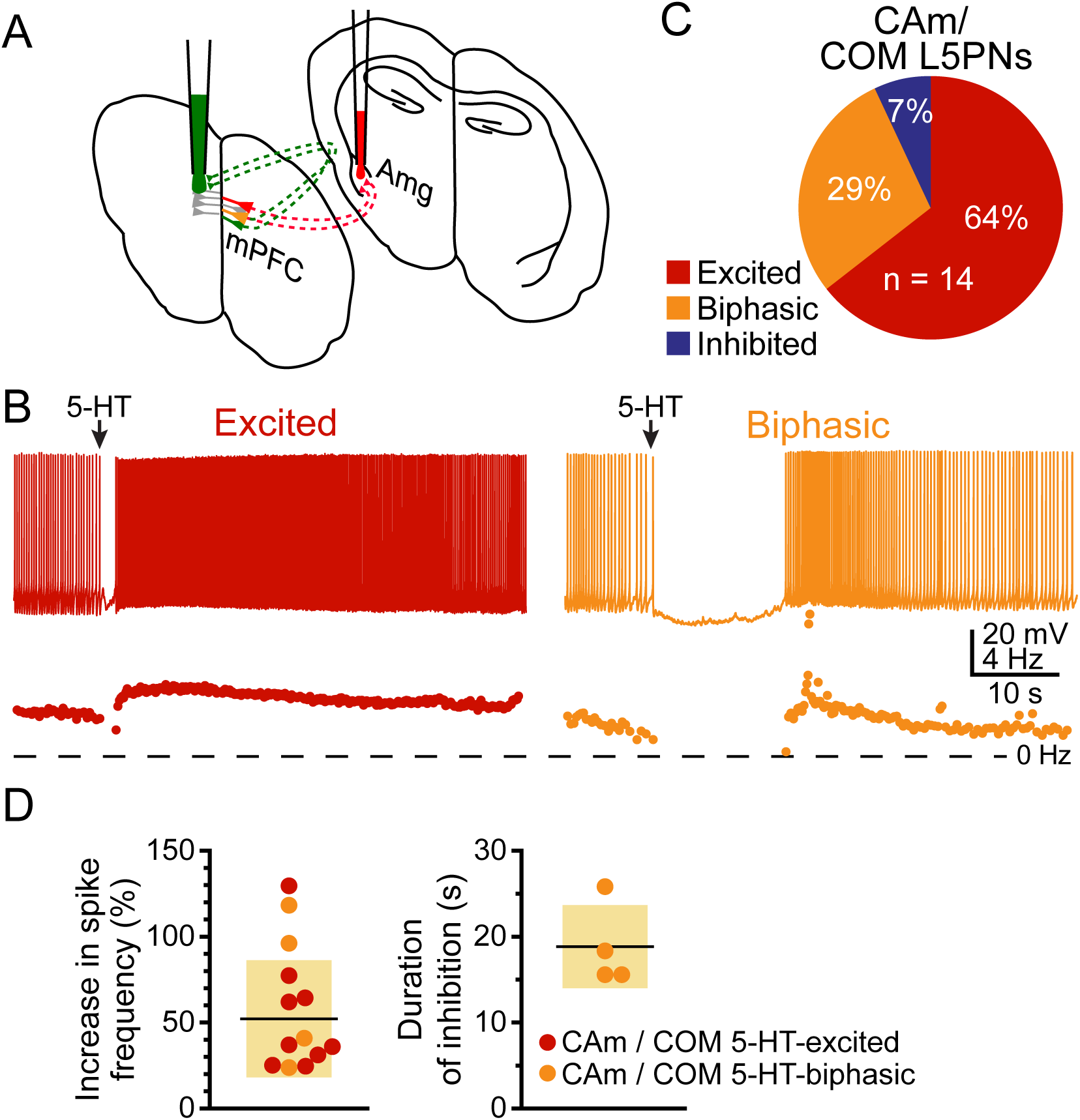
Some CAm neurons are also COM neurons. A. Diagram of dual-labeling of CAm/ COM neurons in the medial prefrontal cortex. **B.** Voltage traces (*top*) and ISF plots (*bottom*) for two double-labeled CAm/COM neurons exhibiting excitatory (red) or biphasic (yellow) responses to 5-HT. **C.** Proportions of 14 CAm/COM double-labeled neurons exhibiting 5-HT responses. **D.** Plots of the magnitudes of 5-HT excitatory responses in 5-HT-excited (red) and 5-HT-biphasic (yellow) CAm/COM neurons (*left*) and the durations of inhibition in 5-HT-biphasic CAm/COM neurons (*right*).

## Discussion

Cortical projections to the amygdala convey top-down control of emotional responses, including fear behaviors (for review, see Courtin et al., 2013). Activity in the prelimbic cortex, likely conveyed to the amygdala via CAm neurons (Stevenson, 2011; Karalis et al., 2016), facilitates emotional responses to conditioned cues, most prominently freezing behavior (Corcoran and Quirk, 2007; Burgos-Robles et al., 2009; Dejean et al., 2016), but also active avoidance (Bravo-Rivera et al., 2014; Bravo-Rivera et al., 2015), suggesting a more generalized role of the prelimbic cortex in goal-directed behaviors and behavioral flexibility (for reviews, see Ragozzino, 2007; Euston et al., 2012; Orsini et al., 2015). Our findings demonstrate that layer 5 CAm neurons in the mouse prelimbic cortex share several morphological and physiological properties with neighboring COM neurons. Specifically, CAm neurons are similar to COM neurons in having sparser dendritic trees, higher input resistances, and less HCN-channel-mediated “sag” potentials than typically observed in neighboring corticofugal (e.g., CPn) neurons. However, HCN-mediated sag potentials in CAm neurons were larger, and RMPs more depolarized, on average, than reported in COM neurons (**Table 1** and **Figure 1C**), suggesting that some differences exist among CAm and COM neurons. Still, we identified double-labeled CAm/COM neurons following dual injections of red and green beads in the amygdala and contralateral cortex, demonstrating that layer 5 CAm neurons overlap to some extent with COM neurons. While we did not test for overlap among layer 2/3 CAm and COM neurons, our findings in layer 5 contrasts with the discrete, non-overlapping, ipsilateral CAm and COM populations reported in layer 2 of the mPFC in Swiss Webster mice (Little and Carter, 2013). This disparity may reflect differences in the laterality of CAm injections (ipsilateral vs contralateral), differences across animal strain, or functional diversity in CAm projections across cortical lamina. However, regardless of population overlap, our study, and that by Little and Carter (2013), agree in finding that CAm and COM neurons are broadly similar in their physiological and morphological properties, including having axonal projections to the contralateral hemisphere.

A main finding of our study is that CAm neurons resemble COM neurons in their responsivity to 5-HT, whereby the vast majority of CAm neurons exhibit 2A-mediated responses, either in the form of purely excitatory responses to 5-HT (in 5-HT-excited neurons), or after transient 1A-mediated inhibition (in 5-HT-biphasic neurons). As with most other physiological measurements, serotonergic responses in CAm neurons were not laterality or sex-dependent (**Table 3**). Further, and as previously revealed in COM neurons (Stephens et al., 2014), 5-HT was unable to excite CAm neurons in the absence of additional excitatory drive, but was sufficient to promote action potential generation in response to suprathreshold simulated synaptic input in 5-HT-excited neurons. This suggests that 5-HT may act to promote cortical output to the amygdala by enhancing the activity of subsets of 2A-expressing CAm neurons that are simultaneously receiving strong excitatory input.

In COM neurons, serotonergic excitation is mediated by G_q_-coupled 2A receptors that engage three postsynaptic ionic effectors to enhance cortical neuron excitability: inhibition of K_V_7 potassium channels and activation of two nonspecific cation conductances (Stephens et al., 2018). Interestingly, these same ionic mechanisms underlie cholinergic excitation of CPn neurons in the prelimbic cortex (Baker et al., 2018), suggesting that they have a conserved role in G_q_-mediated excitation across cortical neuron subtypes and neurotransmitter systems. On the other hand, serotonergic inhibition during biphasic responses results from G_i/o_-coupled 1A receptors enhancing potassium conductances, including those mediated by G-protein-gated inwardly rectifying potassium channels (Andrade and Nicoll, 1987; Luscher et al., 1997). Using specific antagonists, we confirmed that serotonergic excitation and inhibition of CAm neurons are mediated by the same 2A and 1A receptors (respectively) that gate serotonergic responses in COM neurons. While specific ionic mechanisms were not tested in this study, given the overlap in COM and CAm neurons, and the qualitative and quantitative similarities of their 5-HT responses, it would seem likely that a similar set of mechanisms mediates serotonergic responses across both cell types.

### A functional role for serotonergic regulation of CAm neurons

CAm neurons in the prelimbic cortex contribute to reciprocal corticoamygdalar circuits that are engaged during the expression of conditioned fear (Stevenson, 2011; Karalis et al., 2016), and fear conditioning is associated with release of 5-HT into the prelimbic cortex (Hashimoto et al., 1999; Bland et al., 2003). Our results demonstrate that CAm neurons are responsive to 5-HT, suggesting that 5-HT may influence the learning or expression of conditioned fear. Indeed, studies utilizing acute administration of selective serotonergic reuptake inhibitors (SSRIs) to globally increase extracellular 5-HT report enhanced acquisition and expression of cued (but not contextual) fear responses (for reviews, see Bauer, 2015; Bocchio et al., 2016). Further, pharmacological studies have revealed oppositional impact of 1A and 2A receptors in fear expression. For instance, systemic injection of a 2A receptor agonist enhances freezing in response to conditioned cues in rodents (Zhang et al., 2013), while 1A agonists delivered systemically (Inoue et al., 1996; Youn et al., 2009; Ohyama et al., 2016), or locally into the prelimbic cortex (Almada et al., 2015), reduce contextual freezing responses and fear-potentiated startle, respectively. Consistent with a role for 2A receptors in potentiating fear responses, systemic (Zhang et al., 2013) or local blockade of 2A receptors in the prelimbic cortex (León et al., 2017), can impair the expression of conditioned fear.

Since prelimbic CAm neurons exhibit 2A-dependent excitation, 5-HT could contribute to the enhanced functional connectivity observed between the prelimbic cortex and the amygdala during fear behaviors (Karalis et al., 2016). However, the net effect of 5-HT on prelimbic output is difficult to predict because, while 2A receptors may increase the excitability of CAm neurons, 5-HT also influences the excitability of cortical interneurons, and can act presynaptically to regulate glutamate and GABA release (for review, see Puig and Gulledge, 2011). Predictions of 5-HT effects on behavior are further complicated by potentially competing influences of CAm projections arising from different cortical areas (e.g., infralimbic vs prelimbic cortex; see, for instance, Vidal-Gonzalez et al., 2006), and from direct effects of 5-HT on the amygdala or other subcortical structures contributing to fear responses (Bocchio et al., 2016). However, future studies may surmount these limitations by employing genetically targeted approaches that allow for selective manipulation of specific neuron or 5-HT receptor subtypes in restricted brain regions (e.g., prelimbic cortex) at select developmental time points. Such studies, carried out at a systems level, may reveal a role for 5-HT in facilitating not only fear behaviors, but also in regulating an extended intratelencephalic circuit encompassing COM and CAm neurons in both cerebral hemispheres, amygdalar neurons, CA1 neurons in the ventral hippocampus, and their collective overlapping efferents to the striatum (Cho et al., 2013; Heilbronner et al., 2016), to more broadly regulate emotional learning, decision making, and behavior (Vertes, 2006).

## Acknowledgments

The authors thank Saiko Ikeda for technical assistance, and Arielle Baker for discussions.

